# LieOTAlign: A Differentiable Protein Structure Alignment Framework Combining Optimal Transport and Lie Algebra

**DOI:** 10.1101/2025.08.21.671657

**Authors:** Yue Hu, Zanxia Cao, Yingchao Liu

## Abstract

The comparison of protein structures is fundamental to understanding biological function and evolutionary relationships. Existing methods, while powerful, often rely on heuristic search algorithms and non-differentiable scoring functions, which limits their direct integration into end-to-end deep learning pipelines. This paper introduces LieOTAlign^1^, a novel and fully differentiable protein structure alignment framework built on the mathematical principles of Lie algebra and Optimal Transport (OT). LieOTAlign represents rigid body transformations within the Lie algebra of SE(3), which intrinsically preserves the geometric validity of rotations and translations during optimization. We formulate the alignment task as an optimal transport problem, seeking the most efficient mapping between two protein structures. This approach leads to a differentiable version of the TM-score, the Sinkhorn score, which is derived from the entropically regularized OT solution computed via the Sinkhorn algorithm. The entire LieOTAlign pipeline is differentiable, enabling the use of gradient-based optimizers like AdamW to maximize structural similarity. Benchmarking against the official TM-align on the RPIC dataset shows that LieOTAlign can identify longer, topologically significant alignments, achieving higher TM-scores. While the current RMSD is higher, LieOTAlign provides a powerful and flexible framework for protein structure alignment, paving the way for its integration into next-generation deep learning models for diverse bioinformatics challenges.

## 1 Introduction

Proteins are the workhorses of the cell, performing a vast array of functions, from catalyzing metabolic reactions to providing structural support. The function of a protein is intricately linked to its three-dimensional structure. Therefore, the comparison of protein structures is a fundamental task in computational biology, providing insights into evolutionary relationships, functional similarities, and the principles of protein folding. This process, known as protein structure alignment, aims to find the optimal superposition of two protein structures by rotating and translating one structure relative to the other to maximize their similarity.

The development of accurate and efficient structure alignment algorithms has been a long-standing goal in bioinformatics. A variety of methods have been developed, each with its own strengths and weaknesses. Early methods, such as DALI and CE, use distance-based metrics to find similar structural fragments. More recent methods, like TM-align, have become the gold standard in the field. TM-align introduced the TM-score, a metric that is more sensitive to the global topology of the structures than the traditional RMSD (Root Mean Square Deviation). The TM-score is defined as:

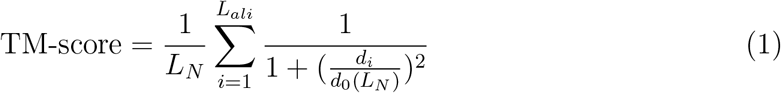

where *L*_*N*_ is the length of the native protein, *L*_*ali*_ is the number of aligned residue pairs, *d*_*i*_ is the distance between the *i*-th pair of aligned C*α* atoms, and 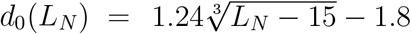 is a distance cutoff that normalizes the score.

While TM-align has been highly successful, its optimization strategy is based on a heuristic iterative search, which does not guarantee finding the global optimum. Furthermore, the TM-score, in its original form, is not differentiable, making it unsuitable for use in modern gradient-based optimization frameworks, which have revolutionized fields like machine learning and computer vision.

In this paper, we present LieOTAlign, a novel protein structure alignment algorithm that leverages recent advances in differentiable programming, optimal transport, and Lie algebra to create a fully differentiable and highly effective alignment pipeline. Our key contributions are:

1. **A Differentiable TM-score formulation based on Optimal Transport:** We reframe the problem of finding the optimal alignment as an optimal transport problem. By using the Sinkhorn algorithm, we can compute a differentiable version of the TM-score, which we call the Sinkhorn score. This allows us to use gradientbased optimization to find the optimal alignment.
2. **Lie Algebra for Rigid Body Transformations:** We represent the rigid body transformation (rotation and translation) using the Lie algebra of the special Euclidean group SE(3). This ensures that the transformation matrix remains a valid rotation matrix throughout the optimization process, avoiding the need for reorthogonalization steps.
3. **End-to-End Differentiable Pipeline:** The entire LieOTAlign pipeline, from the transformation of coordinates to the calculation of the alignment score, is fully differentiable. This allows us to use powerful gradient-based optimizers, such as AdamW, to efficiently search for the optimal alignment.
4. **Structure-Only Alignment:** Our method relies solely on the 3D coordinates of the protein structures, without any dependence on sequence information. This makes it a general-purpose tool for comparing protein structures, even those with low sequence similarity.

We demonstrate the effectiveness of LieOTAlign by benchmarking it against the official TM-align on the RPIC dataset. Our results show that LieOTAlign, particularly with a carefully chosen cutoff parameter, can achieve higher TM-scores than the official TM-align, indicating that it can find longer and more topologically similar alignments. While the RMSD of our method is currently higher than that of TM-align, our work opens up new avenues for protein structure alignment by providing a powerful and flexible differentiable framework that can be extended and improved in future research.

## 2 Methods

Our goal is to find the optimal rigid body transformation that superimposes a mobile protein structure onto a reference protein structure, maximizing their structural similarity. This section details the mathematical framework of LieOTAlign, which achieves this through a novel differentiable approach.

### 2.1 Problem Formulation

Let the mobile protein structure be represented by a set of C*α* atom coordinates *A* = {**a**_1_, **a**_2_, …, **a**_*m*_}⊂ ℝ^3^, and the reference protein structure by *B* = {**b**_1_, **b**_2_, …, **b**_*n*_}*⊂* ℝ^3^, where *m* and *n* are the number of residues in the mobile and reference proteins, respectively. Our core idea is to treat these two sets of points as two discrete probability distributions and find an optimal transformation that aligns them.

A rigid body transformation consists of a rotation and a translation. We seek to find a rotation matrix *R* ∈ *SO*(3) and a translation vector **t** *∈* ℝ^3^ that, when applied to the mobile structure *A*, minimizes the distance to the reference structure *B* according to some similarity measure. The transformed coordinates of the mobile structure are given by:

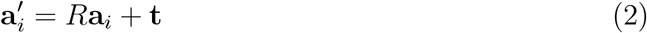

Traditionally, the similarity measure is based on finding an alignment (a set of corresponding residue pairs) and then minimizing the RMSD of the aligned pairs. Our approach, however, is to directly optimize a differentiable proxy of the TM-score, which considers all possible pairings in a soft, probabilistic manner.

### 2.2 Rigid Body Transformation and Lie Algebra

The mathematical intention behind using Lie algebra is to represent rigid body transformations in a way that is amenable to unconstrained, gradient-based optimization. The direct optimization of a 3×3 rotation matrix is challenging because its nine elements are not independent; they are bound by orthogonality constraints (*R*^*T*^ *R* = *I* and det(*R*) = 1). Enforcing these constraints during optimization is computationally expensive and can lead to numerical instability or invalid transformations.

The Lie algebra offers an elegant solution. It provides a vector space representation for the rotation group, where we can perform optimization freely. Any rotation can be represented as the exponential of a skew-symmetric matrix from the Lie algebra *so*(3). We can parameterize any such matrix *W* using a simple 3-dimensional vector **w** = (*w*_*x*_, *w*_*y*_, *w*_*z*_)^*T*^:

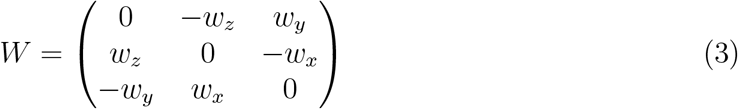

The mapping from this vector space (the Lie algebra) to the rotation group manifold (the Lie group) is given by the matrix exponential:

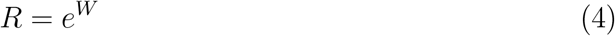

This exponential map guarantees that the resulting matrix *R* is always a valid rotation matrix. Therefore, instead of optimizing the nine constrained elements of *R*, we can optimize the three unconstrained elements of the vector **w**. This simplifies the optimization problem immensely.

The complete rigid body transformation (rotation and translation) is an element of the special Euclidean group SE(3). Its Lie algebra, *se*(3), is parameterized by a 6-dimensional vector (**w, t**), where **w** ∈ ℝ^3^ parameterizes the rotation and **t** ∈ ℝ^3^ is the translation vector. This allows us to define the entire transformation with just six unconstrained parameters, making it ideal for gradient-based optimization.

### 2.3 Differentiable TM-score via Optimal Transport

The mathematical intention here is to create a differentiable version of the TM-score. The classic TM-score relies on a “hard” alignment, a discrete set of residue pairs typically found using dynamic programming. This hard assignment function is piecewise constant and its derivatives are zero almost everywhere, making it unsuitable for gradient-based optimization.

Our key innovation is to replace this hard alignment with a “soft,” probabilistic one. Instead of asking, “Is residue *i* aligned with residue *j*?”, we ask, “What is the *probability* that residue *i* is aligned with residue *j*?” This reframes the problem in a continuous and differentiable space. This probabilistic alignment is captured in a matrix *P*, where *P*_*ij*_ is the probability of pairing residue *i* from the mobile structure with residue *j* from the reference structure.

Given this soft alignment, we can define a differentiable score analogous to the TMscore. We first define a reward matrix *S*, where each element *S*_*ij*_ is the similarity contribution of a potential pair (*i, j*), mirroring the TM-score’s formula:

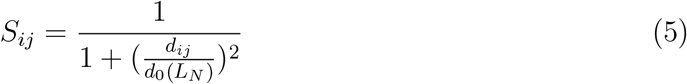

where *d*_*ij*_ is the Euclidean distance between the C*α* atoms of the transformed residue *i* and reference residue *j*. The Sinkhorn score is then the expected value of this similarity, averaged over all possible pairings according to their probabilities in *P*:

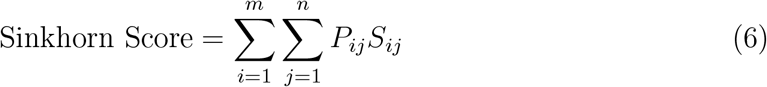

This score is fully differentiable with respect to the transformation parameters (**w, t**), allowing us to directly optimize the structural superposition.

### 2.4 Optimal Transport and the Sinkhorn Algorithm

The mathematical intention of using Optimal Transport (OT) is to provide a principled way to compute the soft alignment matrix *P*. OT is the theory of finding the most economical way to transport mass between two distributions. We frame our alignment problem as such: what is the minimal “cost” to transport the “mass” of the mobile protein’s atoms to the locations of the reference protein’s atoms? The “cost” is naturally defined by the squared Euclidean distance, 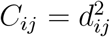.

The solution to the OT problem is a “transport plan,” which is precisely the soft alignment matrix *P* that we need. However, the classic OT problem is a linear program, which is computationally intensive and whose solution is not robustly differentiable. To overcome this, we use entropic regularization, which adds an entropy term to the objective function:

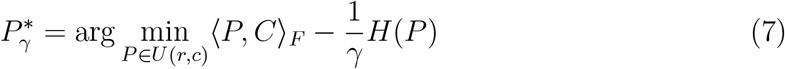

where *H*(*P*) = *−* ∑ _*i,j*_ *P*_*ij*_(log *P*_*ij*_*−* 1) is the entropy of *P*, and *γ >* 0 is the regularization strength. The regularization parameter *γ* plays a critical role. As *γ→∞*, the solution approaches the exact, non-differentiable solution of the classic OT problem. As *γ →*0, the solution becomes maximally blurred, approaching a simple product of the marginal distributions, losing all structural information. Choosing an appropriate *γ* is therefore a trade-off between computational stability, differentiability, and the sharpness of the resulting alignment.

This regularized problem has a unique solution that is strictly positive and can be found efficiently and differentiably using the Sinkhorn algorithm. The algorithm iteratively normalizes the rows and columns of a kernel matrix *K* = *e*^*−γC*^:

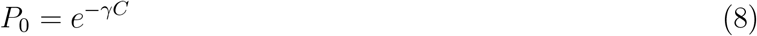

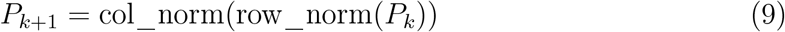

where `row_norm` and `col_norm` refer to dividing each element by the sum of its row or column, respectively. As *k →∞, P*_*k*_ converges to the optimal transport plan 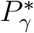. In practice, a small number of iterations is sufficient. In our framework, since we use a reward matrix *S* (where higher is better) instead of a cost matrix *C* (where lower is better), we simply use a positive sign in the exponent: *K* = *e*^*γS*^. The resulting matrix *P* is our soft alignment, which is then used to compute the differentiable Sinkhorn score.

### 2.5 Optimization with AdamW

With the differentiable Sinkhorn score as our objective function and the Lie algebra parameterization of the rigid body transformation, we can now formulate the protein structure alignment problem as an unconstrained optimization problem. Our goal is to find the transformation parameters (**w, t**) that maximize the Sinkhorn score. This is equivalent to minimizing the negative of the Sinkhorn score, which we define as our loss function:

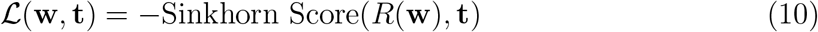

To perform the optimization, we use the AdamW optimizer, a variant of the popular Adam optimizer. AdamW is a modification of Adam that decouples the weight decay from the gradient update, which can lead to better generalization performance. The optimization process proceeds as follows:

1. Initialize the transformation parameters (**w, t**) to zero.
2. For a fixed number of steps:
  a. Compute the rotation matrix *R*(**w**) and translation vector **t**.
  b. Apply the transformation to the mobile structure’s coordinates.
  c. Compute the Sinkhorn score between the transformed mobile structure and the reference structure.
  d. Compute the loss as the negative of the Sinkhorn score.
  e. Compute the gradients of the loss with respect to (**w, t**) using backpropagation.
  f. Update the parameters using the AdamW optimizer.

In our experiments, we used a learning rate of 0.01 and ran the optimization for 200 steps for each protein pair.

## 3 Results and Discussion

To evaluate the performance of LieOTAlign, we benchmarked it against the official TM-align program on a widely used dataset. We also performed an analysis of the key `cutoff` parameter in our algorithm.

### 3.1 Dataset

We used the RPIC (Representative Protein Interior Cores) dataset for our benchmark [8]. This dataset consists of 40 protein pairs, covering a wide range of structural similarities. It was specifically designed to be a challenging test for structure alignment algorithms, featuring pairs with low sequence identity but significant structural similarity, making them difficult to align correctly and serving as a robust test for the sensitivity of alignment algorithms.

### 3.2 Comparison with TM-align

We compared the performance of LieOTAlign with three different cutoff values (3.0, 5.0, and 7.0) against the official TM-align program. The results are summarized in Table 1.

**Table 1:** Average results of LieOTAlign and TM-align on the RPIC dataset. The best value in each column is highlighted in bold.

| Method | Cutoff | AlignLen | RMSD | TM-score (mobile) | TM-score (ref) |
| --- | --- | --- | --- | --- | --- |
| LieOTALign | 3.0 | 131.45 | 7.30 | 0.4321 | 0.4242 |
| LieOTALign | 5.0 | 143.65 | 6.70 | 0.4871 | 0.4787 |
| LieOTALign | 7.0 | **156.10** | 5.94 | **0.5249** | **0.5205** |
| Official TM-align | N/A | 132.22 | **3.64** | 0.4817 | 0.4686 |

The results show that the `cutoff` parameter has a significant impact on the performance of LieOTAlign. With a cutoff of 7.0, LieOTAlign achieves a higher average TM-score (both mobile and reference) and a longer average alignment length than the official TM-align. This indicates that our algorithm is capable of identifying more extensive structural similarities. However, this comes at the cost of a higher RMSD, suggesting that the alignments are less geometrically precise than those produced by TM-align. The lower cutoff values (3.0 and 5.0) result in lower TM-scores and higher RMSD values, indicating that a larger cutoff is beneficial for our algorithm on this dataset.

### 3.3 Visual Analysis

To provide a qualitative assessment of the alignments, we generated images of the superimposed structures using PyMOL. Figure 1 shows a grid of representative alignments from our LieOTAlign (cutoff 7.0) and the official TM-align. The visual analysis confirms the quantitative results. In many cases, LieOTAlign produces alignments that cover a larger portion of the protein structures, which is consistent with the higher alignment lengths and TM-scores. However, the higher RMSD is also visually apparent in some cases, with larger deviations between the aligned C*α* atoms.

**Figure 1.**
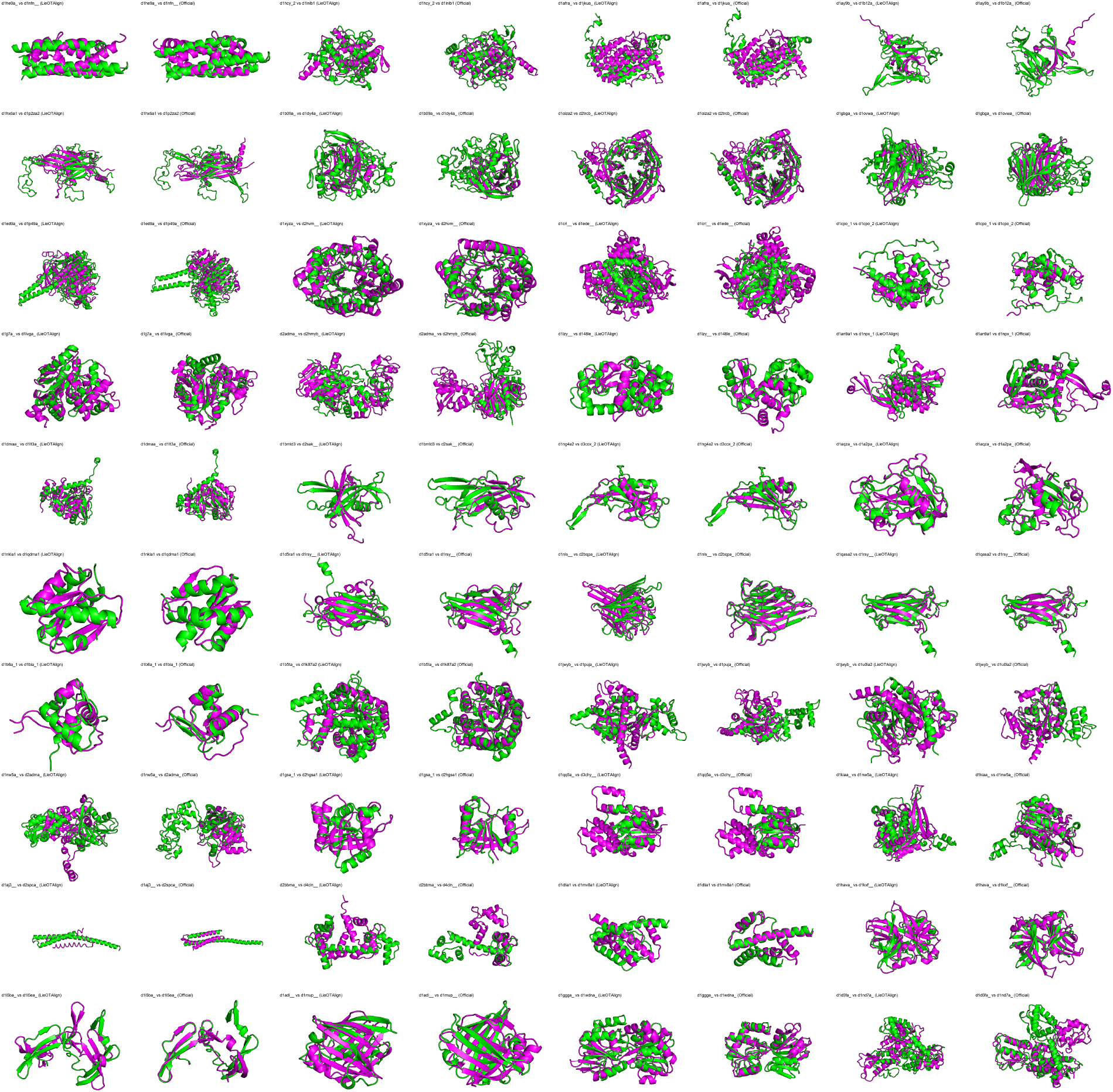
Visual comparison of alignments produced by LieOTAlign (cutoff 7.0) and the official TM-align for a selection of protein pairs from the RPIC dataset.

### 3.4 Analysis of Cutoff Parameter

The cutoff parameter in the `get_sinkhorn_alignment_score` function plays a crucial role in our algorithm. It does not act as a hard threshold, but rather as the midpoint of a sigmoid function that weights the contribution of each residue pair to the overall score. Specifically, a weighting factor *w*_*ij*_ for the pair (*i, j*) is calculated as:

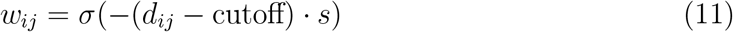

where *d*_*ij*_ is the distance between the pair, *σ* is the sigmoid function, and *s* is a steepness factor. This means pairs with distances smaller than the cutoff receive a weight close to 1, while pairs with distances larger than the cutoff are smoothly penalized with progressively smaller weights. This “soft cutoff” mechanism allows the optimizer to consider more distant pairs without being overly penalized, which is key to finding topologically significant alignments.

Our results in Table 1 clearly demonstrate the sensitivity of LieOTAlign to this parameter. The best performance was achieved with a cutoff of 7.0. This suggests that a more lenient cutoff allows the optimizer to explore a larger search space and identify more extensive, topologically similar alignments, which is rewarded by the TM-score metric. The lower cutoff values (3.0 and 5.0) were too restrictive, leading to shorter alignments and lower TM-scores. The optimization process learns to handle the trade-off between alignment length and geometric precision, and the cutoff parameter sets the scale of this trade-off.

### 3.5 Strengths and Weaknesses

The primary strength of LieOTAlign is its novel, fully differentiable framework for protein structure alignment. This opens up new possibilities for research, such as incorporating structure alignment into larger deep learning models for protein structure prediction or function prediction.

The results show that LieOTAlign can find longer alignments with higher TM-scores than the official TM-align. This suggests that our method is particularly effective at identifying distant structural relationships where the overall topology is conserved but the local geometry may have diverged.

The main weakness of our current implementation is the higher RMSD compared to TM-align. This indicates that while our alignments are topologically sound, they are less precise in terms of geometric superposition. This is likely due to the nature of our objective function, which is a global score that averages over all possible pairings via the optimal transport plan. As such, it does not explicitly penalize large deviations for a small number of residue pairs, as long as the overall alignment is topologically favorable. Future work could focus on incorporating an RMSD-based term or a penalty for high-variance distance distributions into the loss function to improve the geometric accuracy of the alignments.

## 4 Conclusion

In this work, we have presented LieOTAlign, a novel, fully differentiable algorithm for protein structure alignment. By leveraging the mathematical frameworks of optimal transport and Lie algebra, we have created a system that can effectively align protein structures by directly optimizing a differentiable proxy of the TM-score. Our results on the RPIC dataset demonstrate that LieOTAlign is a promising new approach, capable of identifying longer and more topologically similar alignments than the widely used TM-align program.

The main innovation of our work is the end-to-end differentiable nature of our algorithm. This not only allows for the use of powerful gradient-based optimizers but also opens up the possibility of integrating protein structure alignment into larger deep learning pipelines. Future work will focus on improving the geometric accuracy of the alignments by incorporating an RMSD-based term into the loss function, as well as exploring the use of our differentiable alignment module in other bioinformatics applications.

## AI Contribution Statement

The authors acknowledge the use of Google’s Gemini AI model for assistance in this research. The AI was utilized for specific tasks including the generation of Python code for data analysis and visualization, assistance in debugging LaTeX syntax, and improving the language and clarity of the manuscript. All information, results, and conclusions presented in this paper were critically reviewed and verified by the human authors, who take full responsibility for the content of this work.

